# Global mapping of RNA homodimers in living cells

**DOI:** 10.1101/2021.05.13.444021

**Authors:** Marta M Gabryelska, Grzegorz Kudla

## Abstract

RNA homodimerization is important for various physiological processes, including the assembly of membraneless organelles, RNA subcellular localization, and packaging of viral genomes. However, understanding of RNA homodimerization has been hampered by the lack of systematic *in vivo* detection methods. Here we show that PARIS, COMRADES, and other RNA proximity ligation methods can detect RNA homodimers transcriptome-wide as “overlapping” chimeric reads that contain more than one copy of the same sequence. Analysing published proximity ligation datasets, we show that RNA:RNA homodimers mediated by direct base-pairing interactions are rare across the transcriptome, but highly enriched in specific transcripts, including U8 snoRNA, U2 snRNA and a subset of tRNAs. Analysis of data from virus-infected cells reveals homodimerization of SARS-CoV-2 and Zika genomes, mediated by specific palindromic sequences located within protein-coding regions of N protein in SARS-CoV-2 and NS2A gene in Zika. We speculate that regions of viral genomes involved in homodimerization may constitute effective targets for antiviral therapies.

## Introduction

The biological functions of RNA molecules depend on their ability to form intra- and intermolecular interactions, often mediated by Watson-Crick base pairing. Intramolecular base pairing determines the structure and function of RNA, including rRNA and tRNA; it regulates viral replication; and it influences the efficiency of mRNA translation into proteins. Intermolecular RNA-RNA base pairing underlies codon-anticodon recognition, splicing, and regulation of gene expression by miRNA and siRNAs. Intramolecular and intermolecular interactions are interdependent, and according to the competing endogenous RNA (ceRNA) hypothesis (Salmena, et al. 2011; Gardiner, et al. 2015), intermolecular RNA interactions also have the potential to rewire regulatory networks and expand the information encoded in a genome.

A special case of intermolecular interaction between two identical molecules is known as homodimerization. Although homodimers are common in proteins (Bergendahl and Marsh 2017), relatively few homodimers of RNA molecules have been described in vivo (reviewed in (Bou-Nader and Zhang 2020)). Perhaps the best studied are dimers of the HIV genome, which are initiated by an interaction between two copies of the palindromic sequence known as DIS (Berkhout and van Wamel 1996). This interaction leads to the formation of an extended double helix which joins together two copies of the genome, launching a series of events which leads to the packaging of the pair of genomes into one capsid (Paillart, et al. 2004) Homodimerization events have been described in retroviruses, hepatitis C virus, SARS coronavirus, and in bacteriophages (Clever, et al. 2002; Shetty, et al. 2010; Ishimaru, et al. 2013; Dubois, et al. 2018).

RNA oligomerization also plays a role in the process of phase separation, which leads to the formation of membraneless RNA-containing organelles, such as P-bodies, stress granules, nucleoli, Cajal bodies and others (Jain and Vale 2017; Khong, et al. 2017; Nguyen, et al. 2018; Van Treeck and Parker 2018; Van Treeck, et al. 2018). There is growing evidence that such granules are formed via transient protein-RNA and RNA-RNA interactions. As an example, homo- and heterodimerisation of mRNA induces the formation of distinct types of phase-separated droplets in a filamentous fungus (Langdon, et al. 2018). Homodimerization also influences the localization of oskar and bicoid mRNAs in Drosophila embryos (Ferrandon, et al. 1997; Wagner, et al. 2001; Wagner, et al. 2004; Jambor, et al. 2011; Masliah, et al. 2013). Strong interaction between mRNA and pre-mRNA of CUP1 gene leads to RNA miscompartmentalisation and localisation to cytoplasmic foci, possibly including P-bodies and stress granules (Qu, et al. 2014).

An example of pathogenic homodimerization has been observed in a mutated variant of a human mitochondrial tRNA (Wittenhagen and Kelley 2002; Roy, et al. 2005). Additionally, tRNA fragments (tRFs) were shown to form homodimers (Tosar, et al. 2018) and tetramers (Lyons, et al. 2017). CAG and other repeats underlying RNA expansion disorders form hairpin structures, with a stem composed of periodically occurring standard C-G and G-C base pairs (Ciesiolka, et al. 2017). Repeat expansion, corelated with the severity of disorders, increases the possibility of homodimer formation. Sufficiently long tri-nucleotide repeats can form foci in vivo through phase separation (Jain and Vale 2017). Homodimers are also formed by various ribozymes and riboswitches (Bou-Nader and Zhang 2020). Dimerization of RNAs is used in nanobiotechnology for the design and construction of RNA architectures through controlled self-assembly of modular RNA units (tectoRNAs) (Chworos, et al. 2004; Guo 2010; Ishikawa, et al. 2013; Geary, et al. 2014; Grabow and Jaeger 2014; Tanaka, et al. 2016). These observations suggest that transient and stable RNA homodimers play a role in a variety of physiological and pathological processes.

The last few years have seen the development of RNA proximity ligation methods to map cellular RNA-RNA interactions. CLASH (Kudla, et al. 2011), miR-CLIP (Imig, et al. 2015) and hiCLIP (Sugimoto, et al. 2015) use a protein bait to detect protein-associated RNA duplexes, while PARIS (Lu, et al. 2016), LIGR-seq (Sharma, et al. 2016), SPLASH (Aw, et al. 2016) and COMRADES (Ziv, et al. 2018) use a small molecule, psoralen, to crosslink interacting RNA strands. Proximity ligation methods have been commonly used to identify heterotypic interactions, such as interactions between snoRNA, miRNA, piRNA, or sRNA, and their respective targets (Kudla, et al. 2011; Helwak, et al. 2013; Grosswendt, et al. 2014; Ramani, et al. 2015). However, these methods also uncover many homotypic interactions, where the two partners can be mapped to the same gene. Homotypic interactions, usually assumed to originate from the same RNA molecule, have been used to reveal the secondary structures of cellular RNAs (Kudla, et al. 2011; Aw, et al. 2016; Lu, et al. 2016; Sharma, et al. 2016) and structural dynamics of viral genomes (Ziv, et al. 2018; Huber, et al. 2019; Ziv, et al. 2020). However, homotypic interactions could also in principle represent binding between a pair of identical molecules forming an RNA homodimer. Here we describe a method to identify RNA homodimerization events in RNA proximity ligation data by measuring the overlap between arms of chimeric reads. Applying the analysis to published data from multiple proximity ligation experiments, we find that homodimerization is rare in vivo, but is enriched in specific genes and in specific regions of viral genomes.

## Results

### Overlapping chimeras indicate intermolecular interactions

CLASH, PARIS, and other RNA proximity ligation methods rely on the ligation of interacting fragments of RNA, which are then detected as chimeric reads by high-throughput sequencing. We reasoned that chimeras that represent intra- and inter-molecular interactions can be distinguished from each other by an analysis of sequence overlap between the arms of each chimera.

When an RNA molecule that comprises an *intramolecular* interaction is subjected to proximity ligation, the RNA is fragmented into smaller pieces, which - by definition - originate from distinct parts of the RNA. When these fragments are ligated, sequenced, and mapped back to the reference, they should never be mapped to the same region of the RNA. The possible arrangements of the two fragments on the source RNA are shown in Figure 1A. Of these arrangements, the ungapped 5’-3’ chimera is indistinguishable from the source RNA sequence and cannot be identified as a chimera by a simple mapping approach. The other arrangements: gapped 5’-3’, ungapped 3’-5’, and gapped 3’-5’ are all feasible and are commonly detected in proximity ligation experiments.

**Figure 1.**
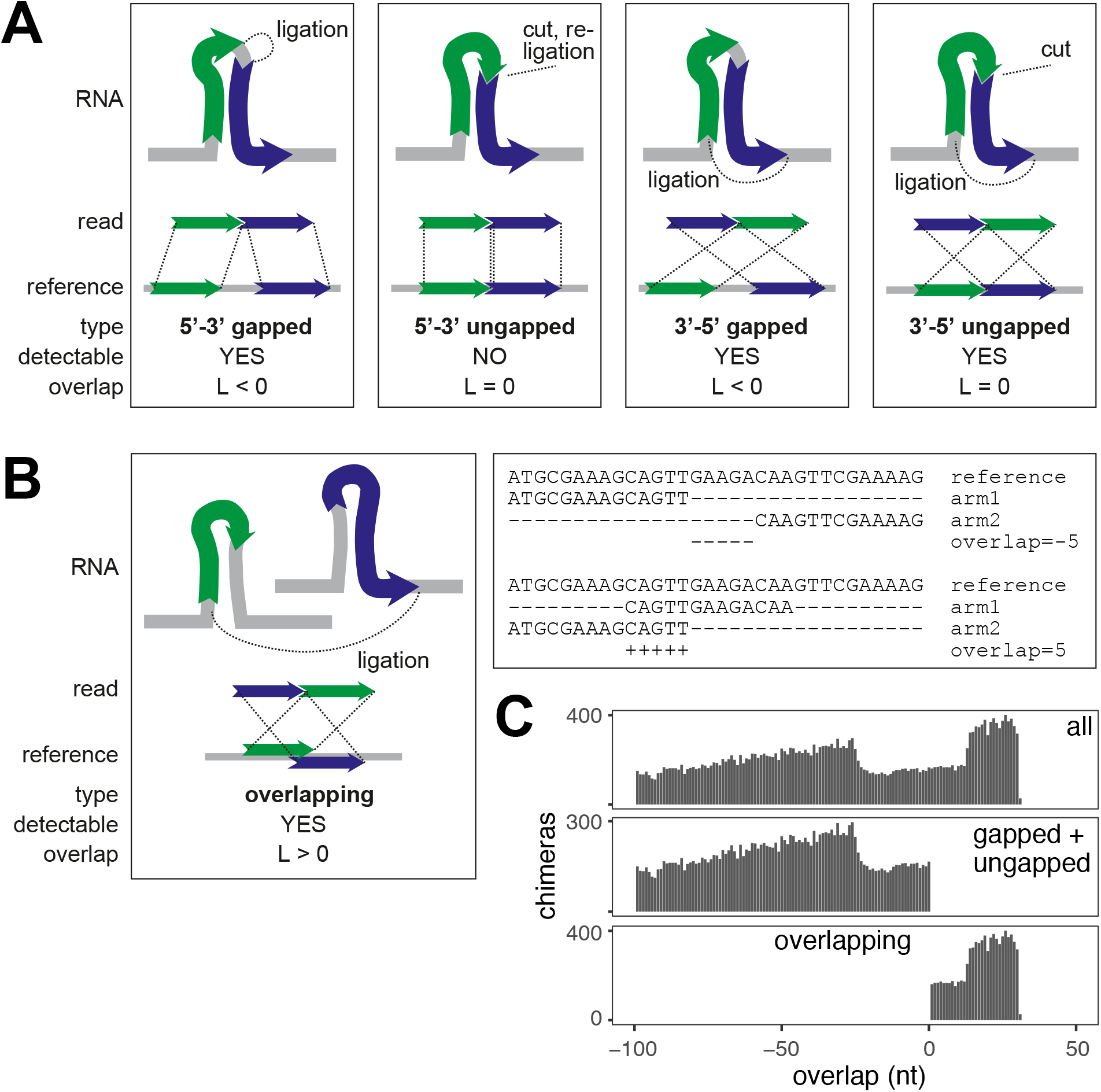
Classification of chimeric reads. (A) Types of non-overlapping chimeras that may be formed by proximity ligation. The RNA fragments, shown as green and blue arrows, can originate from the same transcript, as shown in the figure, or from distinct transcripts (not shown). The dotted lines in the upper panel indicate ligation sites. (B) (left) Diagram of an overlapping chimera. The RNA fragments originate from two distinct copies of a transcript, but are mapped to the same region of the reference gene. (right) Examples of non-overlapping (top) and overlapping (bottom) chimeric reads mapped to a reference gene. (C) Distribution of the calculated overlap score (L) in all simulated chimeras (top), simulated non-overlapping chimeras (middle) and simulating overlapping chimeras (bottom).

In contrast, when a hypothetical *intermolecular* interaction exists between two copies of the same RNA molecule, the two interacting fragments may or may not originate from the same part of the RNA. When these fragments are mapped to the source RNA sequence, they can be found in any of the arrangements shown in Figure 1A, and in an additional overlapping arrangement (Figure 1B). Thus, gapped and ungapped chimeras can result from intra- and inter-molecular interactions, but overlapping chimeras are diagnostic of intermolecular interactions.

In the following sections, we discuss the suitability of bioinformatic methods to detect the relevant types of chimeric reads; the types of chimeras we identify in various RNA proximity ligation experiments; and the possible origin and interpretation of the interactions we detect.

### Detection of overlapping chimeras in simulated sequencing data

To identify chimeras, we used the hyb pipeline (Travis, et al. 2014). Hyb maps reads against a reference sequence database with one of several tools (BLAST, bowtie2, or blat) and detects chimeric reads with two separate local matches in the database. To test whether hyb is suitable for detection of gapped, ungapped, and overlapping chimeras, we assembled a test dataset with simulated chimeras by concatenating all possible pairs of 30-nt substrings from an arbitrary 228-nt RNA sequence. Using either blast or bowtie2 as the mapping engine, hyb correctly identified the majority of sequences as chimeric. A subset of 5’-3’ gapped chimeras and overlapping chimeras were not called by the algorithm (Figure S1). Inspection of the blast and bowtie2 outputs showed that these chimeras were interpreted by the mapping programs as non-chimeric reads with internal deletions or insertions. We also called chimeras with STAR, a general-purpose mapping tool that has been used in some RNA proximity ligation studies. Although the results were comparable with hyb, STAR missed most 5’-3’ chimeras and a subset of 3’-5’ chimeras (Figure S1). Both hyb and STAR commonly mis-identified the position of the ligation junction between the two arms of the chimera by 1-3 nucleotides.

We then quantified the degree of overlap between arms of chimeric reads using a metric, L, defined for chimeras where both arms are mapped to the same reference transcript, as follows:

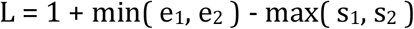

In the equation above, s_1_ and s_2_ are start mapping coordinates of arms 1 and 2 of the chimera on the reference transcript, and e_1_ and e_2_ are end coordinates of arms 1 and 2. L is positive for overlapping chimeras, null for ungapped 3’-5’ chimeras, and negative for gapped 5’-3’ or 3’-5’ chimeras.

We then calculated the distribution of L across chimeras detected in the test dataset. As expected, L was positive for simulated overlapping reads, and negative for simulated non-overlapping reads (Figure 1C). These results show that our methods are appropriate for the identification of overlapping chimeras in RNA proximity ligation data.

### Overlapping chimeras in RNA proximity ligation data

We analysed representative RNA proximity ligation datasets generated by several experimental protocols ((Helwak, et al. 2013; Ramani, et al. 2015; Aw, et al. 2016; Lu, et al. 2016; Sharma, et al. 2016; Waters, et al. 2016; Li, et al. 2018; Ziv, et al. 2018; Huber, et al. 2019; Cai, et al. 2020; Ziv, et al. 2020); see Methods). The protocols differ, among others, in the method employed to stabilise RNA-RNA interactions. CLASH uses UV-protein crosslinking, with only one RNA strand expected to be covalently linked to a protein and the other bound by complementarity. SPLASH, PARIS and COMRADES utilise psoralen crosslinking, while RIC-Seq is based on protein-dependent formaldehyde crosslinking and RPL omits the crosslinking step altogether.

We detected gapped, ungapped and overlapping chimeras in all datasets, but the relative proportions of these chimeras varied greatly between datasets (Figure 2). Strikingly, UV and psoralen crosslinking yielded large numbers of gapped and ungapped, but few overlapping chimeras. Although gapped chimeras could originate from inter- or intra-molecular interactions, the near-absence of overlapping chimeras suggests that homomeric intermolecular interactions are rare in these datasets. By contrast, RPL and RIC-Seq recovered similar numbers of gapped and overlapping chimeras, indicative of RNA homodimerization. Both RPL and RIC-Seq can plausibly recover indirect interactions: RIC-Seq was specifically designed to detect indirect contacts through protein formaldehyde crosslinking, while RPL might allow for reassociation of RNA-RNA complexes during chemical processing *in situ*, due to the absence of a covalent linker. These results suggest that RNA homodimerization mediated by direct RNA-RNA base pairing is uncommon *in vivo*. The results also show that RNA duplexes are generally stable during library preparation, at least in the CLASH, SPLASH, PARIS, and COMRADES methods, because random reassociation of duplexes would lead to the formation of similar proportions of gapped and overlapping chimeras.

**Figure 2.**
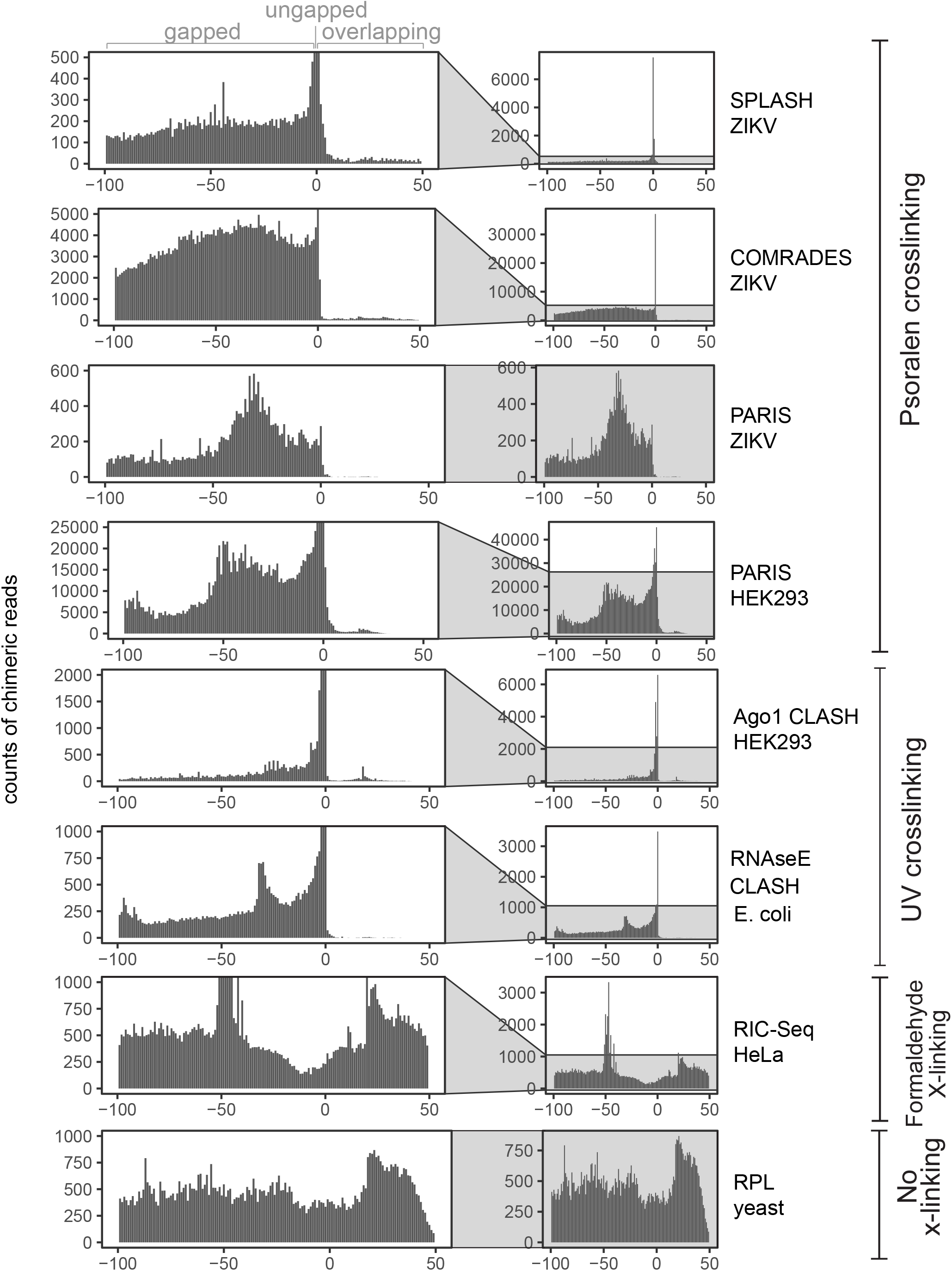
UV- and psoralen-crosslinking predominately generates gapped and ungapped chimeras. The distribution of overlap scores (L) across proximity ligation datasets: SPLASH ZIKV (Zika virus) (Huber, et al. 2019); COMRADES ZIKV (Ziv, et al. 2018); PARIS ZIKV (Li, et al. 2018); PARIS HEK293 (human) (Lu, et al. 2016); Ago1 CLASH HEK293 (Helwak, et al. 2013); RNAse E CLASH (*Escherichia coli*) (Waters, et al. 2016); RIC-Seq HeLa (human) (Cai, et al. 2020); RPL (*Saccharomyces cerevisiae*) (Ramani, et al. 2015) (see Methods).

An intriguing pattern is the peak at L=0 in Figure 2, indicating the preferential recovery of 3’-5’ ungapped chimeras relative to gapped chimeras (as discussed above, 5’-3’ ungapped chimeras cannot be detected with our methods). We propose that ungapped chimeras typically arise from local RNA stem-loop structures, which are subject to three endonucleolytic cleavages, followed by ligation of the distal ends to each other, while gapped chimeras could be created either by four independent endonucleolytic events, or by a combination of three endonucleolytic cuts, combined with exonucleolytic trimming of RNA ends. While enrichment of ungapped chimeras can be readily explained for intramolecular interactions, it is difficult to imagine a mechanism that could enrich ungapped chimeras for intermolecular interactions. These results reinforce our conclusion that stable RNA homodimers are rarely formed in vivo.

### Homodimerization of human and yeast RNAs

Although few RNA homodimers were present in UV- and psoralen-crosslinking experiments, we hypothesized that homodimers might be limited to a specific subset of RNAs. To investigate this possibility, we analysed chimeras detected in individual genes in transcriptome-wide PARIS data from HEK293 cells. To increase the stringency of our analysis, we filtered the data to remove likely mapping errors, chimeras with thermodynamically unstable interactions, homopolymers, and chimeras with very short overlaps (less than 5 nucleotides) (Figure S2). After filtering, gapped chimeras were more common overall, but 47 genes contained overlapping chimeras (Table S1), including 3 genes that were significantly enriched in overlapping chimeras (Fisher’s exact test with Benjamini-Hochberg correction, p<0.05). The most highly enriched transcript was mRNA TMEM107, which contained 30-times more overlapping than gapped chimeras (Figure 3).

**Figure 3.**
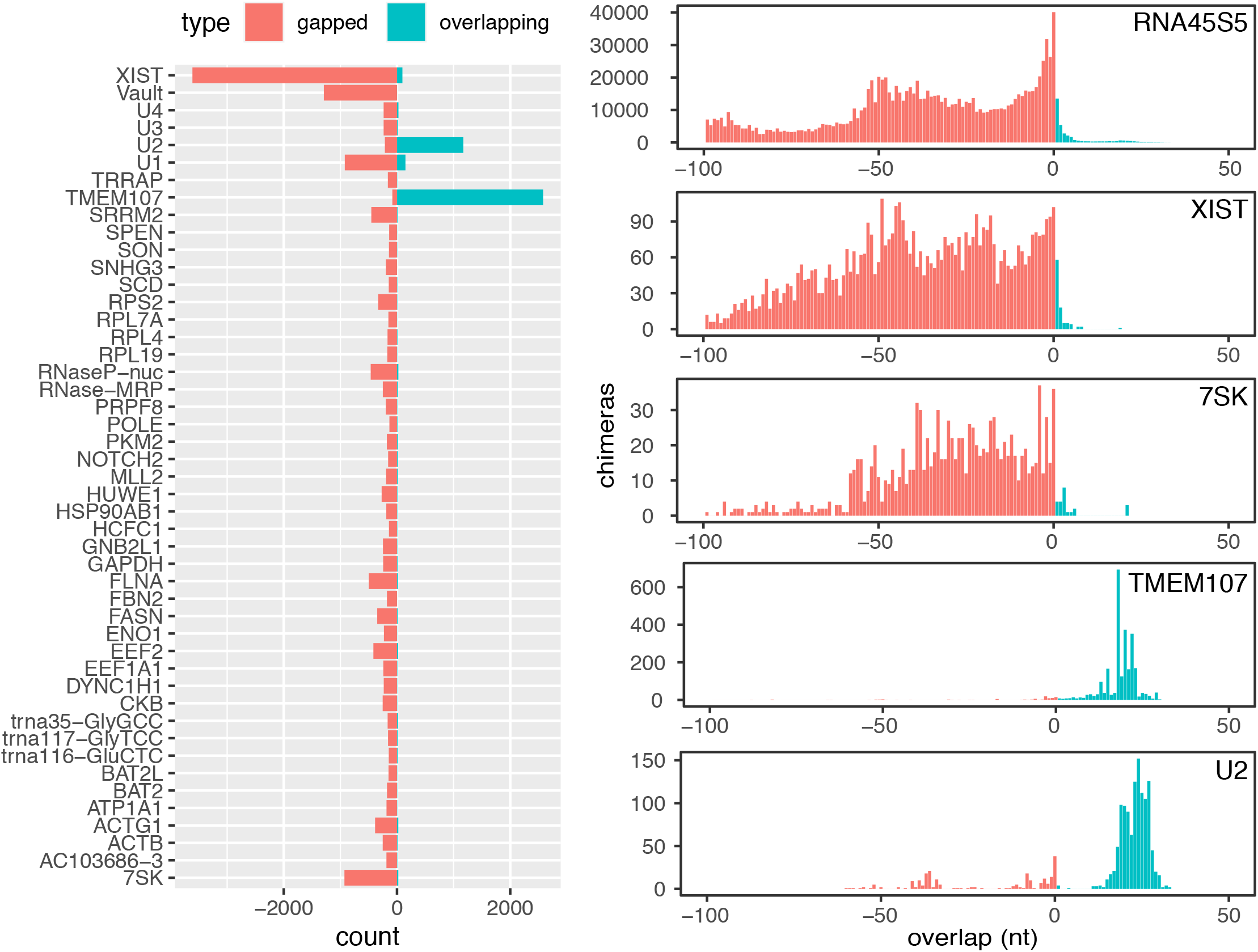
Distribution of overlap scores in individual genes in PARIS experiment. Counts of gapped and ungapped chimeras (pink) and overlapping chimeras (blue) in individual genes in PARIS data from human HEK293 cells (Lu, et al. 2016) (left), and distribution of the overlap value L in selected genes.

TMEM107 contains a small nucleolar RNA (snoRNA), U8, in its 3’ untranslated region, and almost all TMEM107:TMEM107 chimeras mapped to that region, suggesting that these chimeras represent U8:U8 interactions. The chimeras were concentrated around the 5’ end of U8 (Figure 4), and RNA folding prediction showed extended self-complementarity in this part of the transcript, consistent with homodimerization with a predicted free energy of −21 kcal/mol. Previous studies showed that the 5’ region of U8 may base-pair with pre-ribosomal RNA (Peculis 1997), and with the 3’ end of a 3’-extended precursor of U8 (Badrock, et al. 2020). As homodimerization seems incompatible with these interactions, it might represent an immature form of U8, or play a role in the regulation of U8 function. This is potentially important for the pathogenesis of LCC, a neurological disease caused by loss-of-function mutations in U8 (Jenkinson, et al. 2016; Badrock, et al. 2020). Interestingly, some overlapping chimeras comprised a 5’-extended form of U8, indicating that the homodimers may be formed during snoRNA maturation. We have not found any interactions involving other regions of the TMEM107 transcript (Figure S3), nor have we found homodimers of other mRNAs from the TMEM family.

**Figure 4.**
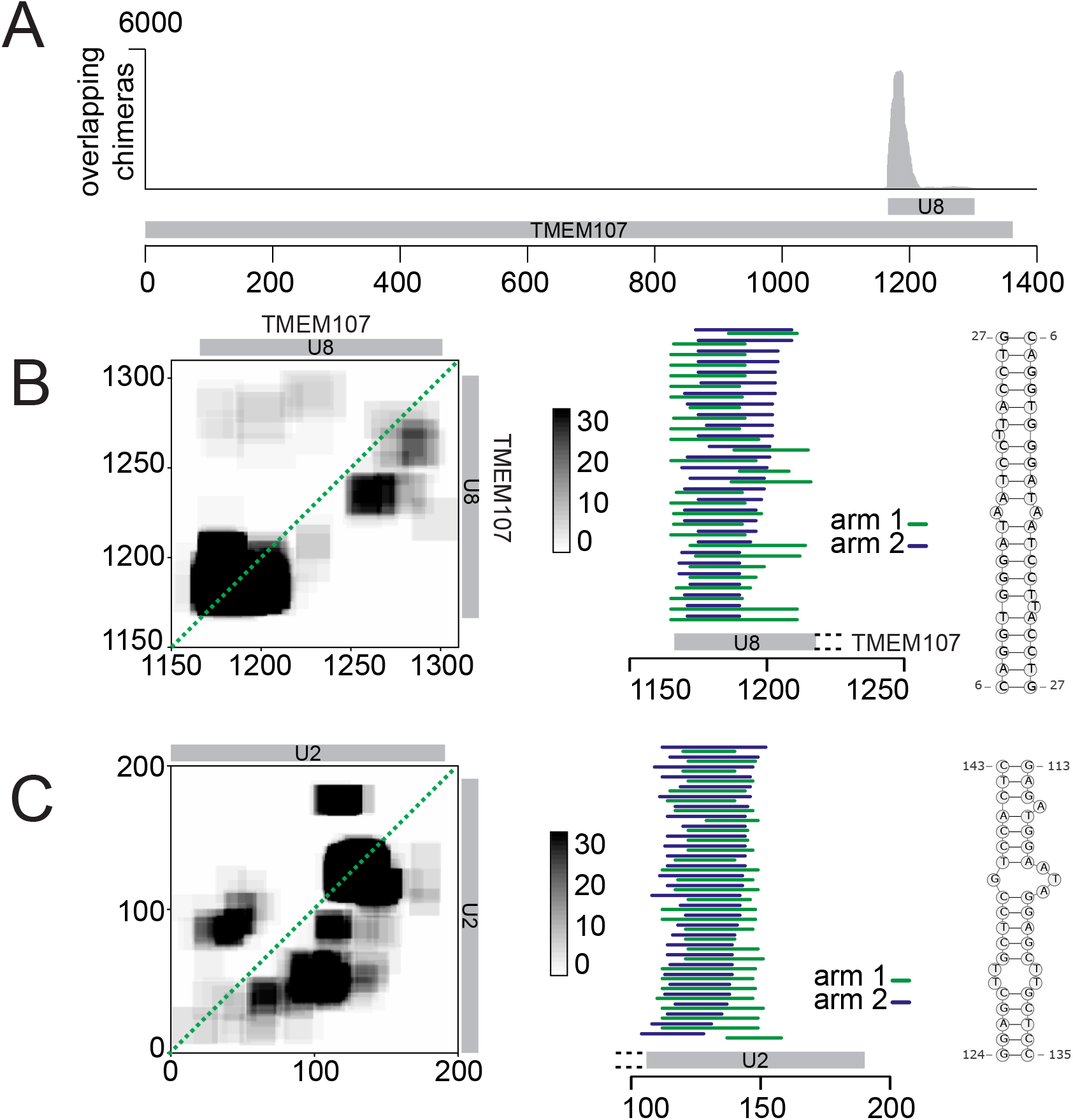
Homodimerization of U8 snoRNA and U2 snRNA. (A) Distribution of overlapping chimeras along the TMEM107 gene in PARIS (HEK293 cells)(Lu, et al. 2016). All overlapping chimeras coincide with the position of U8 snoRNA in the 3’UTR of TMEM107 transcript. (B) (Left) Coverage map of U8:U8 chimeras. Numbers on the X and Y axes indicate positions in the TMEM107 transcript. The position of mature U8 snoRNA is indicated by the grey bars. (Middle) mapping positions of both arms of chimeras along TMEM107. Some chimeras extend beyond the 5’ end of mature U8 (grey bar), indicating that they originate from the U8 precursor molecule. (Right) Predicted base pairing of U8:U8 homdimer. (C) As above, analysis of U2:U2 chimeras. Coordinates indicate positions along the U2 transcript.

In addition to U8:U8 interactions, analysis of PARIS data showed enrichment of homodimers in U1 and U2 snRNA. U2 snRNA contained 4-5 times as many overlapping as gapped chimeras. Regions involved in homodimeric interactions in U1 and U2 are limited to a particular fragment of the RNA, while other types of interactions can be found along the transcript (Figure S3). Most overlapping chimeras in U2 included the sequence of stem loop III, downstream of the Sm binding site, suggesting that in a fraction of U2 molecules found in the cell, stem loop III is unfolded and forms homomeric intermolecular interactions (Figure 4). Out of the two major isoforms of U2, U2-1 shows more efficient deposition of the Sm ring and incorporation into snRNP complexes than U2-2 (Kosmyna, et al. 2020), and we hypothesized that failure of assembly into an snRNP complex might be associated with U2:U2 dimerization. However, analysis of the exact sequences of U2:U2 overlapping chimeras showed that out of 223 reads that could be assigned to specific U2 isoforms, 162 were of the U2-1:U2-1 type, and 59 were U2-1:U2-2 chimeras, suggesting that both U2 isoforms may form homo- and heteromeric intermolecular interactions. U2 homodimers were also found in RIC-Seq and CLASH datasets.

We then compared specific homodimerization events detected by different proximity ligation methods. PARIS and COMRADES showed the largest fractions of homotypic chimeras (Figure S4A), most of which were non-overlapping and likely represented intramolecular interactions. The largest ratio of overlapping to homotypic chimeras was recovered by RIC-Seq (7%, Fig S4A). RIC-Seq also recovered the highest number of genes with overlapping chimeras (over a thousand), 17 of which were significantly enriched for overlapping chimeras (Fig S4, Table S3). Across all RNA biotypes, mRNA formed most homodimers, particularly in RIC-seq, PARIS, and COMRADES, but tRNAs were enriched for homodimers in Ago1-CLASH and SPLASH (Figure S4, S5). tRNA-derived small RNAs (tsRNAs), including tRNA-derived fragments (tRFs) and tRNA halves (tiRNAs), are small regulatory RNAs processed from mature tRNAs or precursor tRNAs (Xie, et al. 2020). tX(XXX)D, a yeast tRNA similar to serine tRNAs (Chan and Lowe 2009) formed a homodimer through a 12 base pair long stem in SPLASH data (Figure S6). The tRNA homodimers detected by Ago1-CLASH in human cells (Figure S6) may indicate a miRNA-tRNA network resulting in competition for binding sites and availability for gene silencing, as reported previously (Shigematsu and Kirino 2015). Interestingly, RIC-Seq showed significant enrichment of homodimers in 20 mitochondrial mRNAs, with CO1, ND2 and ND4 containing the highest numbers of homodimeric reads (Fig S5, S7). Bidirectional transcription of mitochondrial RNA is known to result in hybridization of complementary strands (Dhir, et al. 2018; Kim, et al. 2018), but in the homodimers found in RIC-Seq data, both partners come from the same strand, suggesting that they represent a distinct type of interaction. The mitochondrial mRNA:mRNA interactions showed low thermodynamic stability and short regions of complementarity (2 to 8 base pairs), as expected for indirect interactions facilitated by the high local concentrations of these transcripts in mitochondria. Other homodimerization examples include SNORD12, enriched in overlapping chimeras in COMRADES (Fig S7), and YLR154W-E, a possible ncRNA from yeast with a strong enrichment in homodimers in the RPL data, predicted to dimerize through an extended stem structure (Fig S6).

### Homodimerization of virus RNA

We next turned to COMRADES data from cells that have been infected with SARS-CoV-2 and Zika viruses, to detect possible homodimers of virus RNA. Although Zika RNA is not known to homodimerize, dimerization is an essential step in the packaging of some viruses, including HIV, while dimerization of SARS-CoV RNA was suggested to play a role in translational frameshifting (Ishimaru, et al. 2013). To detect dimers of virus RNA, we analysed the coverage of overlapping chimeras along viral genomes. Unlike gapped chimeras, which covered the Zika genome relatively evenly, overlapping chimeras were strongly enriched in several positions within the NS2A, NS2B and NS5 coding sequences of the Zika virus, indicating possible dimerization sites (Figure 5). RNA folding prediction showed regions of self-complementarity in the interaction sites, including a pair of uninterrupted 11-bp duplexes in the (3578-3656):(3578-3656) region in the NS2A gene. However, folding energy alone was not enough to predict dimerization sites, as evidenced by the weak negative correlation between the count of overlapping chimeras in a genomic window and the predicted strength of homodimeric interaction in that window (pearson R=−0.17, p=3×10^−8^).

**Figure 5.**
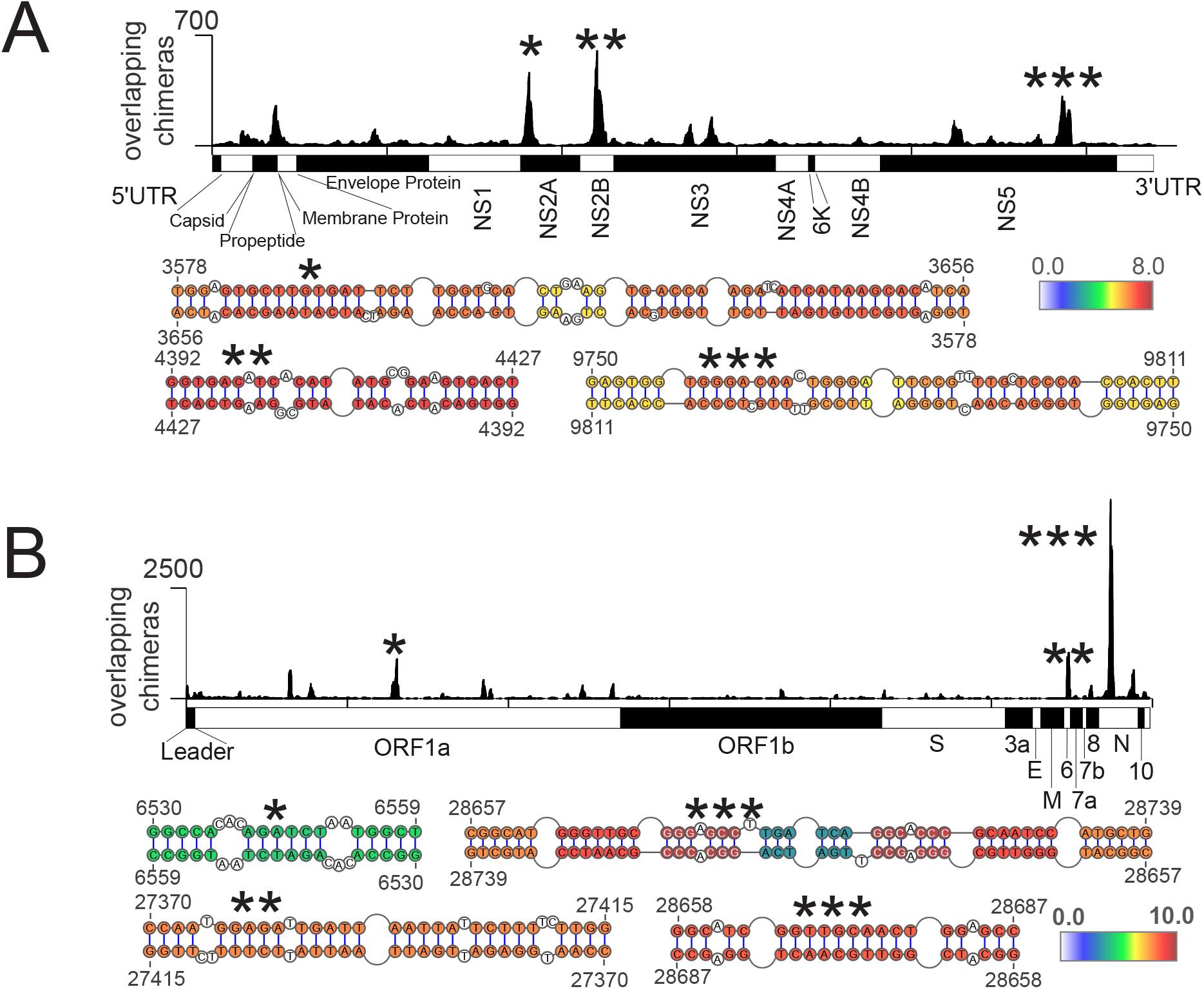
Homodimers of viral genomes. (A) (top) Coverage of overlapping chimeras in Zika virus identified by COMRADES (Ziv, et al. 2018). The regions with the largest coverage of homodimers are indicated by asterisks. (bottom) Predicted secondary structures of regions with high coverage of overlapping chimeras. The colours indicate the COMRADES score, i.e. the log2 of the number of chimeric reads supporting each base pair. (B) Coverage of overlapping chimeras along the SARS-CoV-2 genome identified by COMRADES (Ziv, et al. 2020). The top homodimeric regions and their predicted structures are shown, as described above.

**Figure 6.**
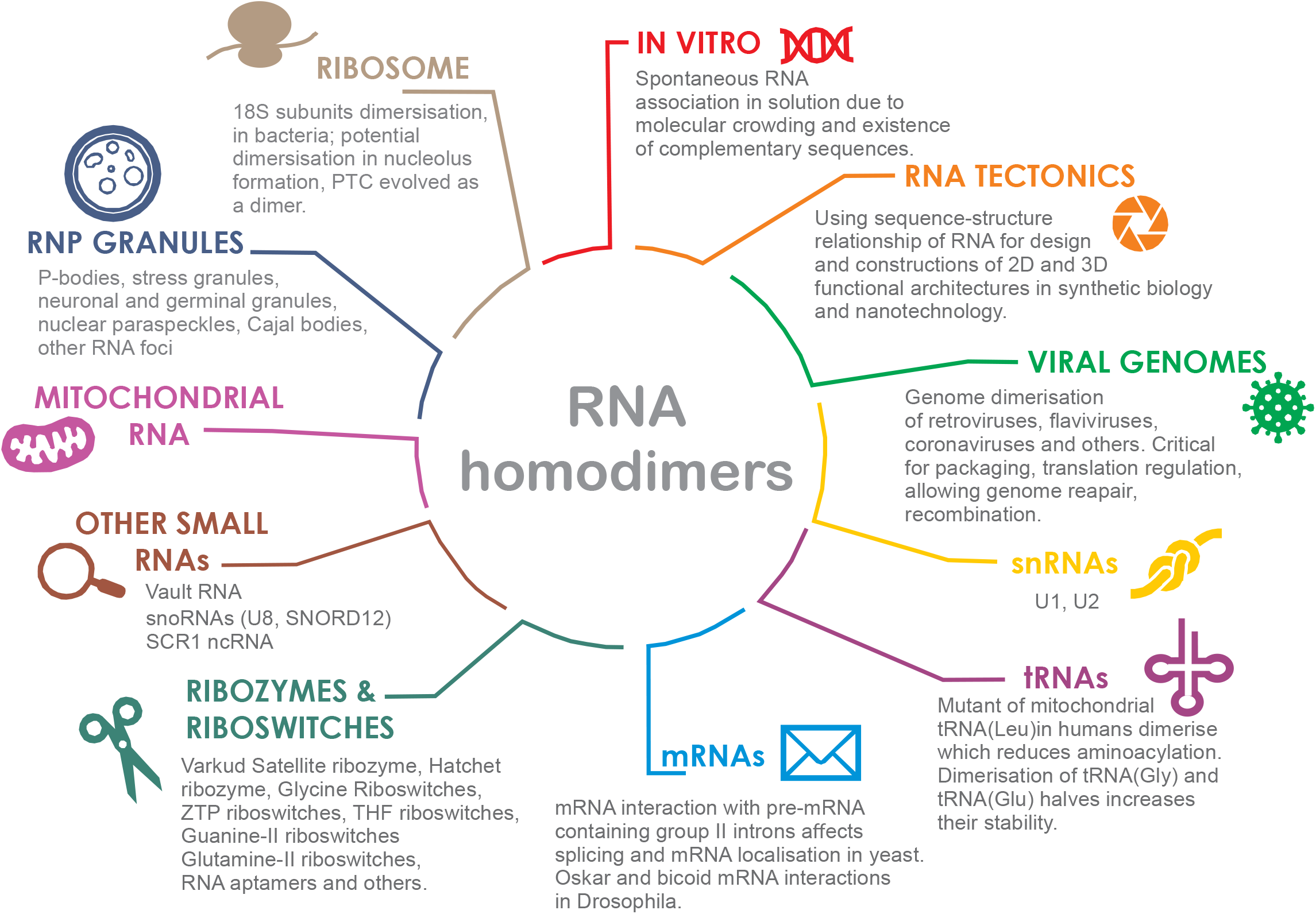
Summary of known and newly identified RNA-RNA homodimers. (Micklem, et al. 2000; Wittenhagen and Kelley 2002; Irion and St Johnston 2007; Ishikawa, et al. 2013; Qu, et al. 2014; Franken, et al. 2017; Dubois, et al. 2018; Tosar, et al. 2018; Van Treeck and Parker 2018; Bou-Nader and Zhang 2020).

We also detected dimerization events in the SARS-CoV-2 genome, with the largest peak in the nucleprotein (N protein) gene, and additional peaks in the N protein, Orf 6 and Orf 1a coding sequences (Figure 5). The region with the largest coverage of overlapping chimeras was 200-nt long (coordinates 28,610-28,810) and the resolution was insufficient to indicate the exact base pairing, but RNA folding analysis showed several high-scoring interactions, including a 10-nt duplex formed by the palindromic sequence, GGTTGCAACT. Although a previous NMR study detected a functionally important dimerization site near the frameshifting element of the SARS-CoV virus (Ishimaru, et al. 2013), our analysis shows no obvious enrichment of overlapping chimeras in the homologous region of SARS-CoV-2.

## Discussion

Although homo-oligomerization is common in proteins, few RNA homo-oligomers have been described in vivo. This is somewhat surprising, given that RNA molecules readily homodimerize in vitro, to the point that special procedures have to be used to isolate monomeric forms of certain RNAs for structural studies (Zhang and Ferre-D’Amare 2014; Bou-Nader and Zhang 2020). The paucity of in vivo homo-oligomers might be explained by the folding of RNAs and by their association with protein complexes, which reduce the propensity for trans-RNA-RNA interactions. Alternatively, the apparent lack of in vivo homodimers might simply reflect the lack of systematic studies of dimerization. Here, by analysing the relative proportions of gapped, ungapped and overlapping chimeric reads in RNA proximity ligation experiments, we find that homodimerization mediated by direct RNA basepairing is indeed rare in vivo. Surprisingly, however, we find that certain human RNAs and some regions of the RNA genomes of the Zika and SARS-CoV-2 viruses are enriched for in vivo homodimers.

Out of thousands of RNAs we examined, only a handful show clear evidence of dimerization. The propensity to dimerize is necessarily influenced by the primary sequence of the RNA: for example, palindromic sequences or CAG repeats might be prone to form intermolecular interactions. A recent review of RNA homodimer structures detected in viruses, ribozymes, and riboswitches identified preferences for certain sequence and structural arrangements, such as palindromes, complementary strand swapping and kissing-loop interactions (Bou-Nader and Zhang 2020). Indeed, in the present study, palindromic sequences were found in several RNA homodimers. Homodimerization is also likely to be influenced by folding kinetics, association with other proteins and RNAs, subcellular localisation and local concentration of RNA and metal ions. RNA molecules which fold co-transcriptionally into stable secondary structures are unlikely to form extended duplexes with other RNAs (Yu, et al. 2021), whereas molecules that are unfolded by helicases, or located in granules with high local concentrations of a given RNA, might be more likely to form transient or stable oligomers. Copies of RNA molecules located in close proximity may initially interact with a few nucleotides, followed by destabilisation of local structure and nucleation of longer interactions (Ganser, et al. 2019).

Are the RNA homodimers detected by proximity ligation biologically relevant, or experimental artefacts? We argue that non-specific dimerization and oligomerization of RNA during library preparation, if present, should lead to the formation of many overlapping chimeras, distributed across a large variety of RNAs. Indeed, this is what we observe in the RPL and RIC-Seq datasets. RPL is performed without crosslinking, while RIC-seq involves formaldehyde crosslinking. As a result, overlapping chimeras detected by these methods likely indicate local transcript proximity rather than direct basepairing, though it is also possible that a fraction of overlapping chimeras arises during the library preparation step.

By contrast, techniques that rely on UV or psoralen crosslinking: CLASH, SPLASH, PARIS, and COMRADES, are expected to detect RNA-RNA contacts mediated by direct base pairing. We observed that these methods generate few overlapping chimeras, but these chimeras are strongly enriched in a small subset of RNAs, suggestive of bona-fide interactions. Alternatively, overlapping reads might theoretically arise through reverse transcription of an endogenous circular RNA (circRNA), or of an artificial circRNA created in vitro by ligation, producing a concatemeric cDNA. However, the low abundance of circRNAs and low efficiency of RNA ligases makes such events unlikely. We also note that proximity ligation can only identify a subset of possible RNA homodimers, namely those where both RNAs interact via the same part of their sequence, or via two regions that are close enough in the primary sequence to detect overlaps in chimeric reads. Although many known RNA homodimers are of this type (for example, the DIS-DIS’ interaction in HIV, the SL2-SL2 interaction in Moloney murine sarcoma virus (MoMuSV), or the dimerization of Oskar RNA via its 3’ UTR in Drosophila embryos) (Berkhout and van Wamel 1996; Kim and Tinoco 2000; Jambor, et al. 2011), interactions mediated via distant fragments of RNA would not be detectable by proximity ligation.

Homodimerization has now been reported for most major biotypes of RNA, and known roles of homodimers include the packaging of viral genomes, assembly of membraneless organelles, regulation of RNA localization (Figure 5B). Given its dependence on the local concentration of RNA, dimerization might play a role in RNA quorum sensing - a process analogous to that used by bacteria and viruses to coordinate their behaviour in response to the local population density. Nevertheless, many RNA homodimers do not have a known biological function, and indeed might be detrimental. Stretches of dsRNA are known to trigger antiviral immunity through PKR and other cellular factors [Hull and Bevilacqua 2016], and some types of homodimers might be misidentified as foreign RNA. RNA multimerization has also been associated with general cellular stress [Van Treeck and Parker 2018, Van Treeck et al 2018]. In RNA proximity ligation methods, the use of psoralen, formaldehyde and UV light is a stress factor that might contribute to RNA multimerization. In any case, further functional studies are required to elucidate the roles of the wide variety of RNA homodimers that can be detected in our cells.

## Methods

### Benchmarking chimera detection on test data

To benchmark methods for detection of overlapping chimeras, we assembled a test dataset using an arbitrary RNA sequence (nucleotides 1-228 of S. cerevisiae RDN37 gene: Genbank Sequence ID: CP026300.1, range 448071 to 448298, minus strand). We generated all 30-nt substrings of the reference sequence and concatenated all possible pairs of substrings, which yielded 10,871 overlapping chimeras and 28,730 non-overlapping chimeras.

We then called chimeras in the overlapping and non-overlapping datasets using hyb and STAR, using the following commands:

hyb (bowtie2 mapping):

hyb analyse in=input.fasta db=RDN37 format=comp eval=0.001

hyb (blast mapping)

hyb analyse in=input.fasta db=RDN37 format=comp align=blastall eval=0.001

STAR:

STAR --genomeDir. --readFilesIn input.fasta --outFileNamePrefix 06 -- outReadsUnmapped Fastx --outFilterMismatchNoverLmax 0.05 -- outFilterMatchNmin 16 --outFilterScoreMinOverLread 0 -- outFilterMatchNminOverLread 0 --clip3pAdapterMMp 0.1 –chimSegmentMin 15 --scoreGapNoncan -4 --scoreGapATAC -4 --chimJunctionOverhangMin 15

We used the ua.hyb files from hyb and Chimeric.out.junction files from STAR for downstream analysis. To generate the coverage heatmaps of chimeras detected in the test dataset, we extracted the coordinates of chimera junctions and plotted them using Java TreeView (Saldanha 2004).

### Calculation of chimera overlaps

To quantify overlap between arms of chimeric reads, we defined the overlap metric, L, as:

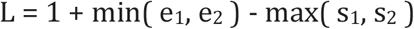

L is defined for chimeras where both arms are mapped to the same reference transcript, or same chromosome in case of mapping to a genome reference. e_1_ represents the end mapping coordinate of the left arm of the chimera (arm 1); e_2_ - the end mapping coordinate of the right arm of the chimera (arm 2); s_1_ and s_2_ represent the start mapping coordinates of the respective arms. Calculation of L was implemented as a custom awk script, taking the ua.hyb files produced by the hyb pipeline as inputs.

### Experimental data

We downloaded the data from the following sources: *Escherichia coli* RNAse E CLASH: GSE77463 (Waters, et al. 2016); human Ago1 CLASH: GSE50452 (Helwak, et al. 2013); human SPLASH: SRR3404931 (Aw, et al. 2016); Zika SPLASH: SRR6252011 (Huber, et al. 2019); Zika COMRADES: E-MTAB-6427 (Ziv, et al. 2018); human LIGR-Seq: SRR3361013 (Sharma, et al. 2016); human PARIS: SRR2814765 (Lu, et al. 2016); Zika PARIS: PRJEB28648 (Li, et al. 2018); human RIC-Seq: SRR8632820 (Cai, et al. 2020); *Saccharomyces cerevisiae* RPL: SRR2048219 (Ramani, et al. 2015); SARS-CoV-2 COMRADES: GSM4676632 (Ziv, et al. 2020).

Sequencing data were downloaded in fastq format (except for the SARS-CoV-2 dataset, where hyb output files were downloaded). Chimeric reads were called and annotated with the hyb package (Travis, et al. 2014) with default settings, using the appropriate transcriptome database (Helwak, et al. 2013; Waters, et al. 2016; Ziv, et al. 2018), as described in Table S4.

Overlap statistics across experimental datasets were visualized in R using the ggplot2 and ggforce libraries (facet_zoom function). To identify genes enriched in overlapping chimeras, we filtered hyb outputs to remove possible mapping errors (any reads with mono- or dinucleotide repeats of length 15 or more, and chimeras where either arm had a mapping e-value greater than 0.0001); we also removed chimeras with predicted interaction energy weaker than −5 kcal/mol. Because very short overlaps might represent mapping or sequencing errors, we conservatively called chimeras with overlap score L>5 as overlapping, and, for consistency, chimeras with −50<L<−5 as non-overlapping. We then assembled a contingency table with counts of overlapping and non-overlapping chimeras for the focal gene and for all other genes, and used a Fisher’s exact test with Benjamini-Hochberg multiple testing correction to identify genes with significant enrichment of overlapping chimeras.

---

**Figure S1.**
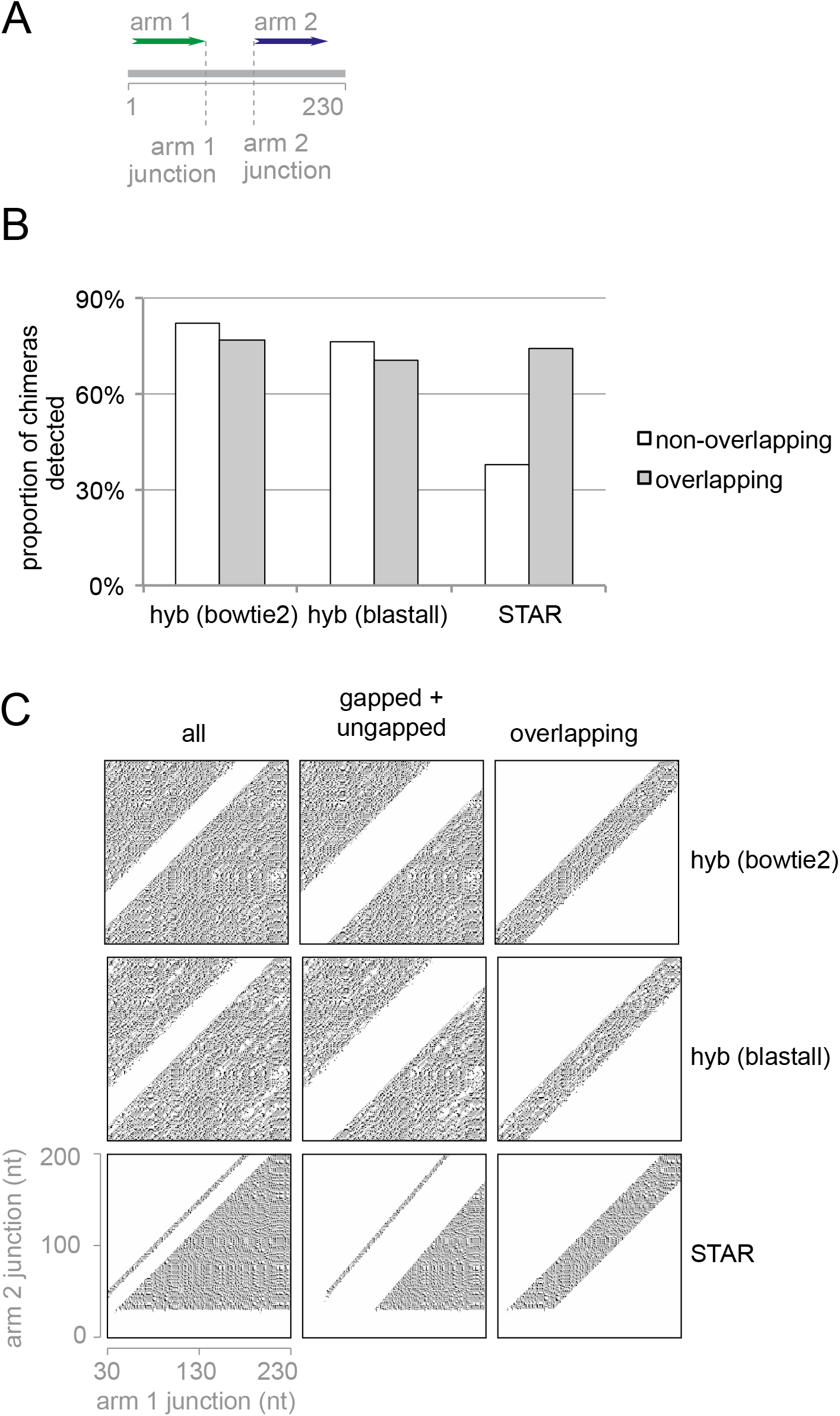
Benchmarking of chimera mapping tools on simulated chimeras. (A) Design of benchmarking dataset: all pairs of 30-nt substrings of a 228-nt long yeast RNA sequence were concatenated, separated into overlapping and non-overlapping sets, and their junction positions were recorded. (B) Proportion of simulated chimeras correctly identified as chimeric by hyb and STAR. (C) Coverage heatmap of chimeric junction positions identified in the test dataset by hyb and STAR.

**Figure S2.**
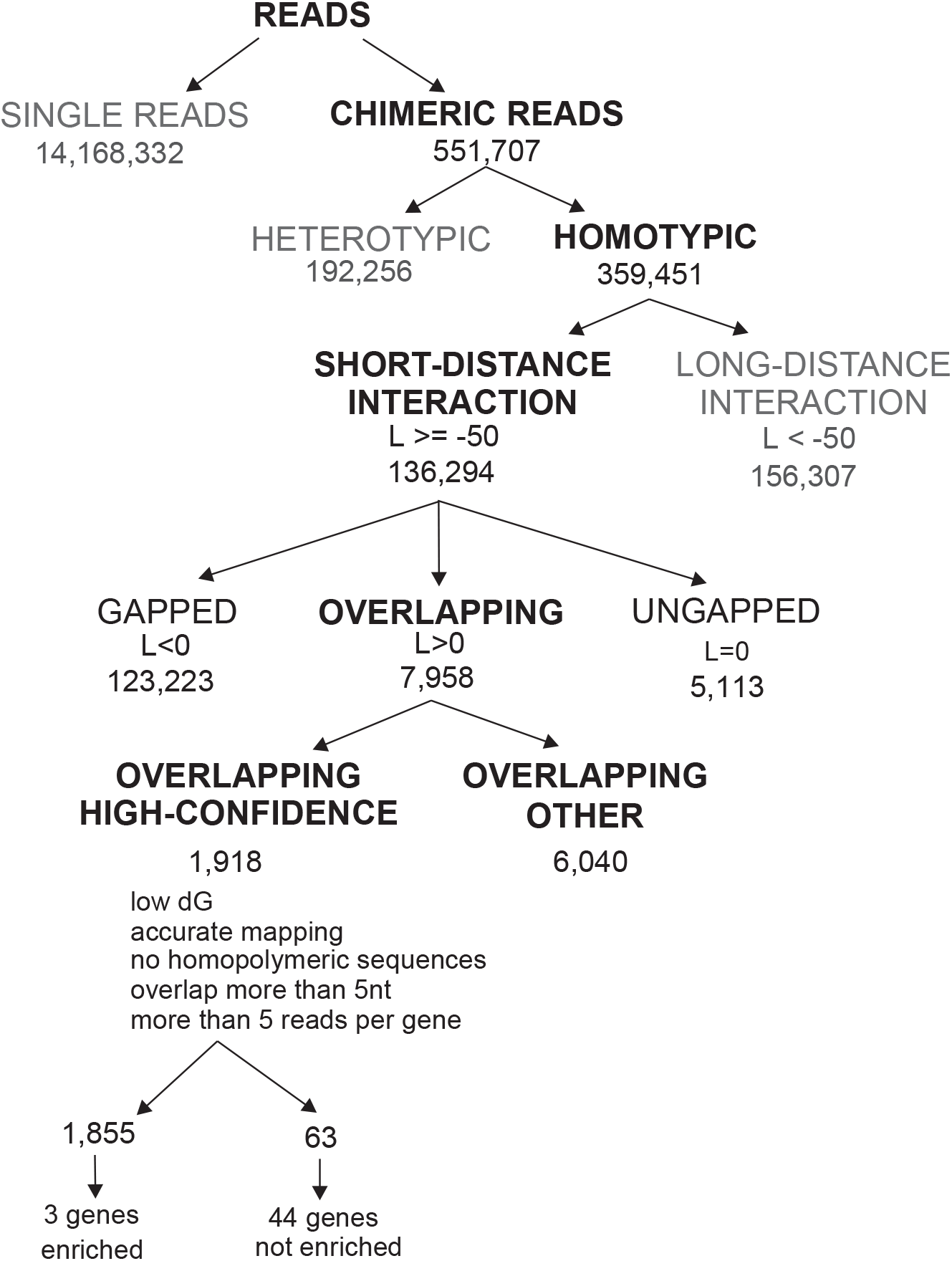
Statistics of chimeric reads found in HEK293 PARIS (Lu, et al. 2016)

**Figure S3.**
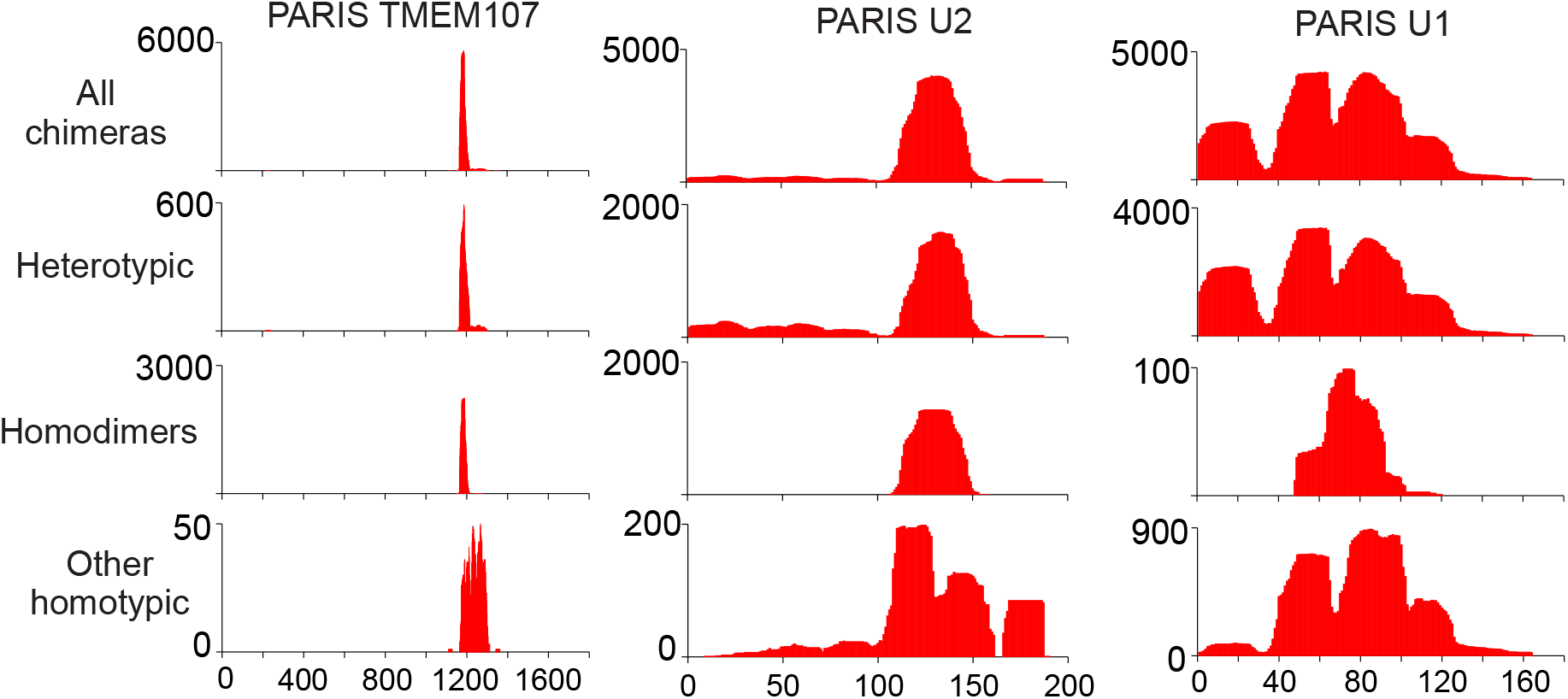
Genes enriched in homodimeric reads in HEK293 PARIS (Lu, et al. 2016). The X axis represents the position along the indicated gene, and the Y axis, chimera coverage. From top to bottom, the graphs represent, for each gene, the coverage of all chimeras, heterotypic chimeras, homotypic overlapping chimeras (homodimers), and other homotypic chimeras.

**Figure S4.**
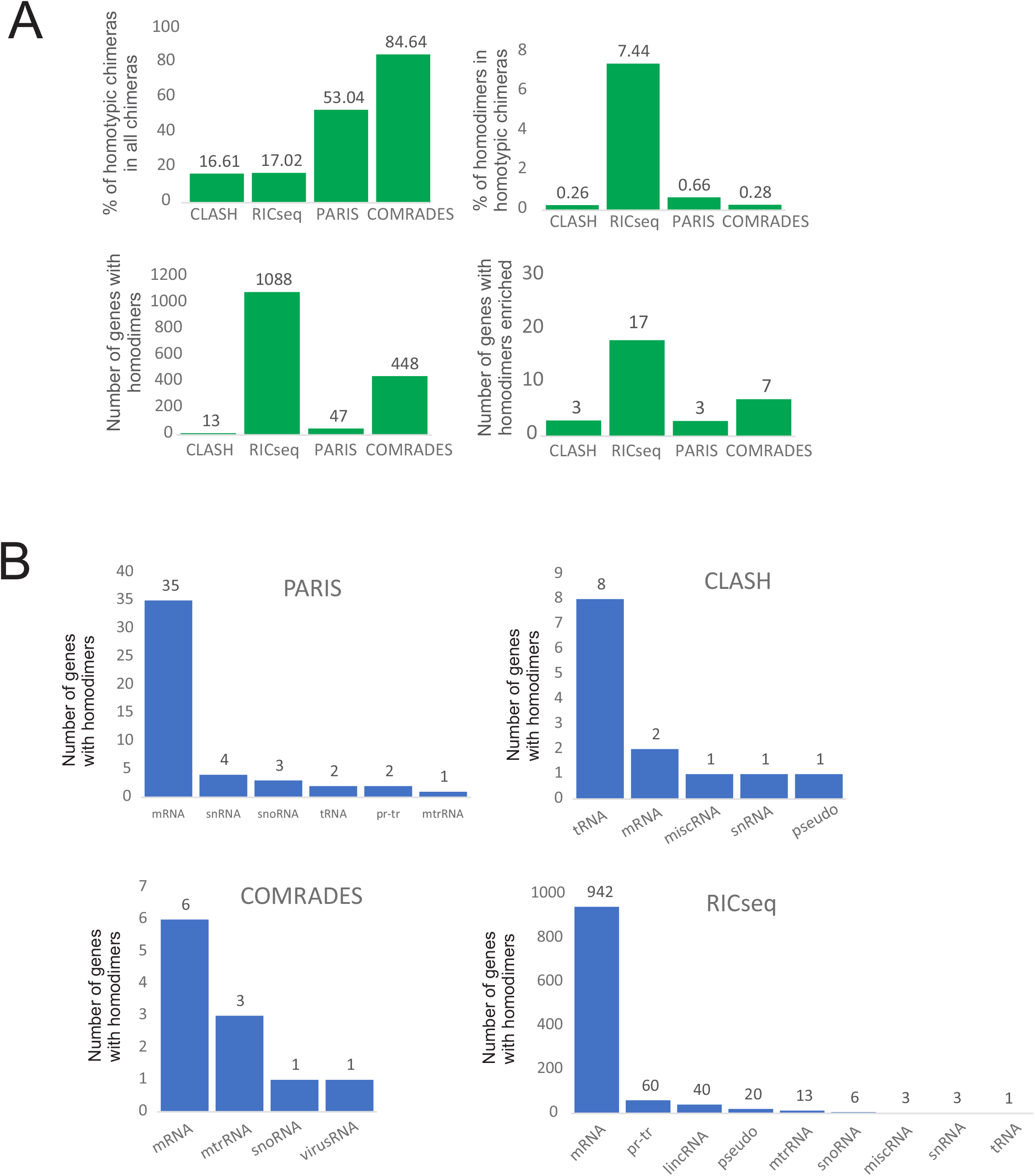
Statistics of homodimers in CLASH (Helwak, et al. 2013), RIC-seq (Cai, et al. 2020), COMRADES (Ziv, et al. 2018) and PARIS (Lu, et al. 2016) data. (A) (top left) Percentage of homotypic chimeras among all chimeras; (top right) percentage of homodimers (overlapping chimeras) among homotypic chimeras; (bottom) Number of genes containing homodimeric reads (left) and genes statistically enriched in homodimeric reads. (B) Classification of genes with homodimers by RNA biotype.

**Figure S5.**
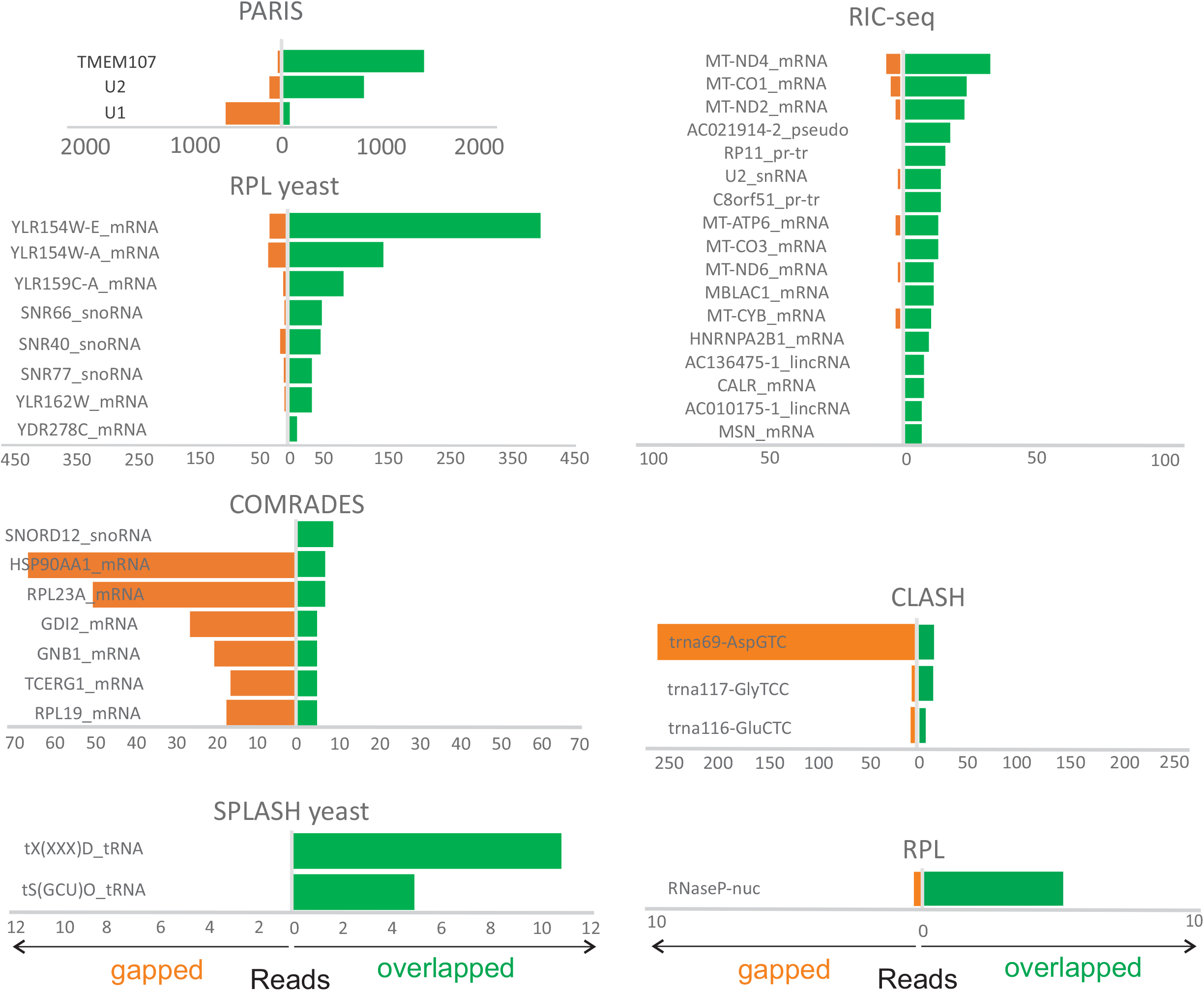
Genes with significantly enriched homodimers in various proximity ligation experiments. Counts of gapped and ungapped chimeras (orange) and overlapping chimeras (green) in individual genes in human PARIS data (Lu, et al. 2016); yeast RPL (Ramani, et al. 2015) human RIC-Seq (Cai, et al. 2020); Zika COMRADES (Ziv, et al. 2018); human Ago1 CLASH (Helwak, et al. 2013); yeast SPLASH (Aw, et al. 2016) and human RPL (Ramani, et al. 2015).

**Figure S6.**
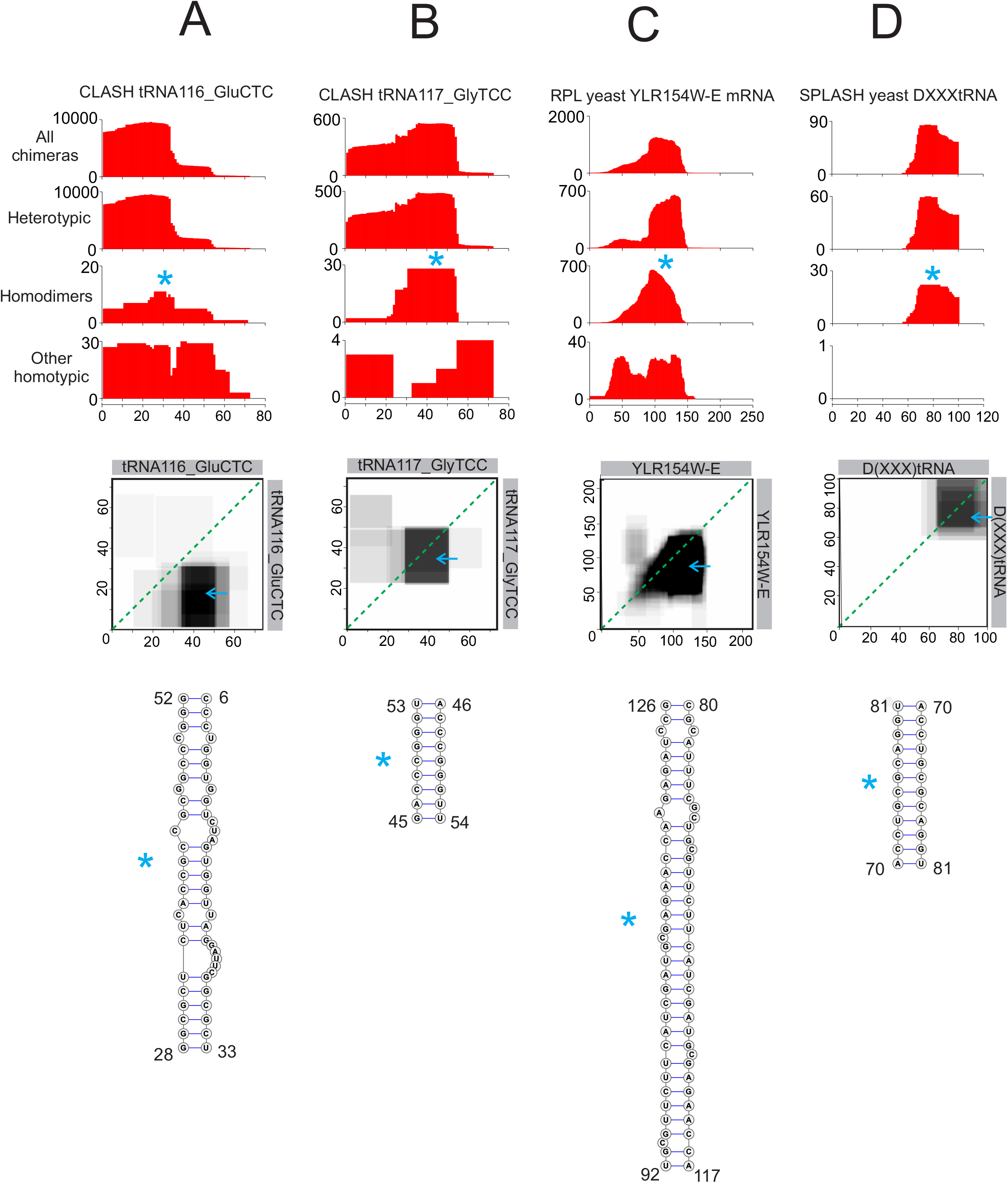
Example homodimers detected by CLASH (Helwak, et al. 2013), RPL (Ramani, et al. 2015), and SPLASH (Aw, et al. 2016).

**Figure S7.**
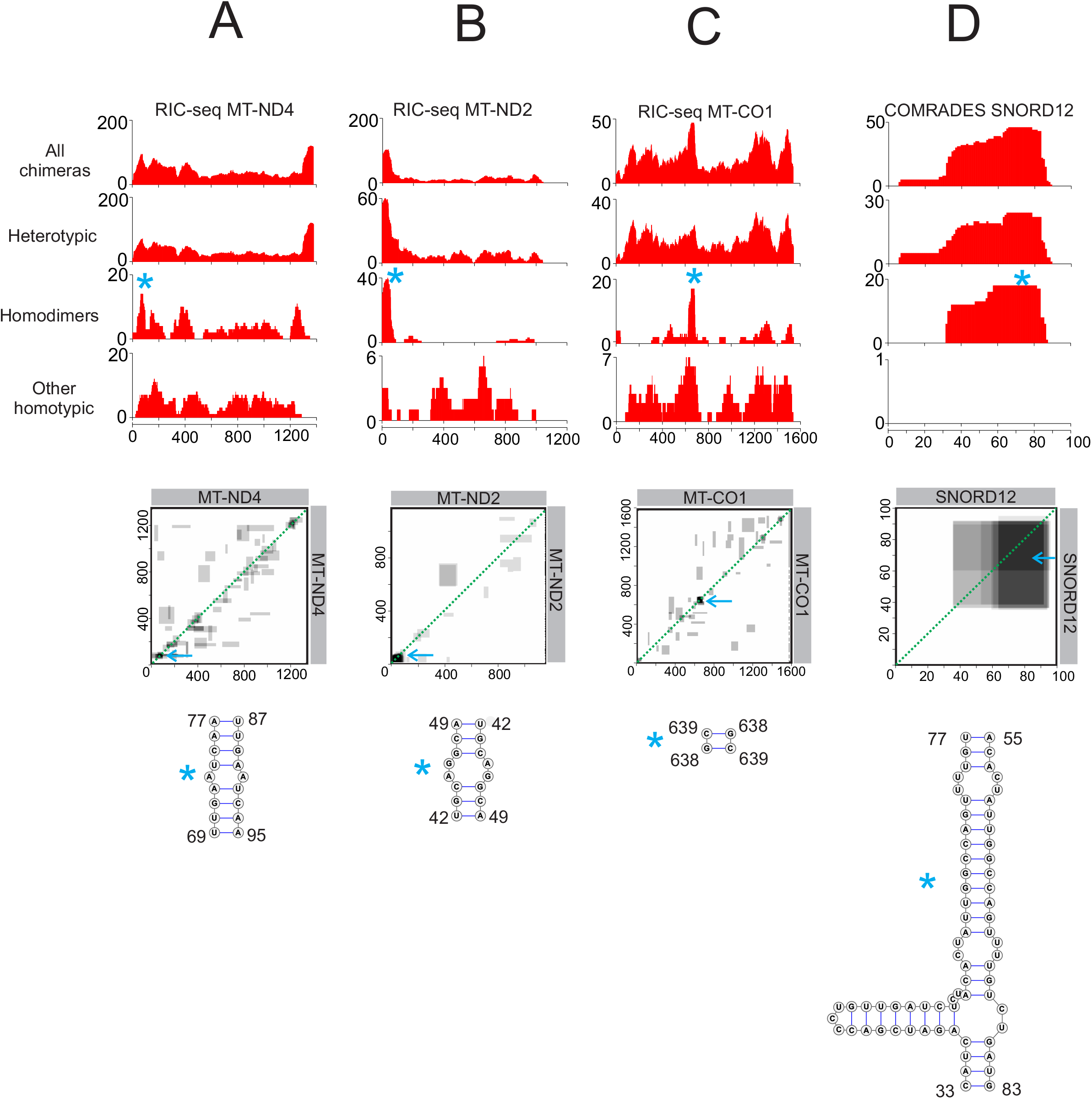
Example homodimers detected by RIC-Seq (Cai, et al. 2020) and COMRADES (Ziv, et al. 2018)

